# An automated BIDS-App for brain segmentation of human fetal functional MRI data

**DOI:** 10.1101/2022.09.02.506391

**Authors:** Emily S. Nichols, Susana Correa, Peter Van Dyken, Jason Kai, Tristan Kuehn, Sandrine de Ribaupierre, Emma G. Duerden, Ali R. Khan

**Author notes:** These authors contributed equally (Emma G. Duerden, Ali R. Khan). Corresponding author: Emily S. Nichols, PhD, Applied Psychology, Faculty of Education, Room 1131, 1137 Western Rd London, Ontario N6G 1G7.

## Abstract

Fetal functional magnetic resonance imaging (fMRI) offers critical insight into the developing brain and could aid in predicting developmental outcomes. As the fetal brain is surrounded by heterogeneous tissue, it is not possible to use adult- or child-based segmentation toolboxes. Manually-segmented masks can be used to extract the fetal brain; however, this comes at significant time costs. Here, we present a new BIDS App for masking fetal fMRI, *funcmasker-flex*, that overcomes these issues with a robust 3D convolutional neural network (U-net) architecture implemented in an extensible and transparent Snakemake workflow. Open-access fetal fMRI data with manual brain masks from 159 fetuses (1103 total volumes) were used for training and testing the U-net model. We also tested generalizability of the model using 82 locally acquired functional scans from 19 fetuses, which included over 2300 manually segmented volumes. Dice metrics were used to compare performance of *funcmasker-flex* to the ground truth manually segmented volumes, and segmentations were consistently robust (all Dice metrics ≥0.74). The tool is freely available and can be applied to any BIDS dataset containing fetal bold sequences. *funcmasker-flex* reduces the need for manual segmentation, even when applied to novel fetal functional datasets, resulting in significant time-cost savings for performing fetal fMRI analysis.

## 1. Introduction

Over the last 2 decades, research in fetal MRI protocols has increasingly been used to non-invasively study the functional, metabolic and structural origins of the fetal brain *in vivo* (Huisman et al., 2002; Prayer et al., 2004; Rousseau et al., 2006; Thomason et al., 2013, 2015; Wheelock et al., 2019). Improved methods to study the fetal brain *in vivo* makes it possible to understand not only how typical, healthy brain development occurs and predicts later cognitive and motor outcomes, but also allows us to study atypical development (Arroyo et al., 2019; Rajagopalan et al., 2021). As fetal functional MRI is becoming more common, there has been an increase in demand for automatic preprocessing and analysis software; while there is an abundance of neuroimaging analysis software for infant, child, and adults, there is currently a dearth of fetus-specific tools, making the process of preprocessing fetal neuroimaging data, especially functional MRI, quite difficult and time-consuming.

One of the most complicated steps in the fetal preprocessing pipeline is brain extraction from the echo-planar imaging (EPI) sequences, the process of isolating the brain within the image, creating a “mask” and stripping away the skull and surrounding tissue laying outside of the identified region. In pediatric and adult samples this step is straightforward; numerous tools have been developed, with algorithms relying heavily on identifying the dark cerebrospinal fluid that separates the brain from the skull (Kalavathi & Prasath, 2016). As the fetal brain does not have clear boundaries separating it from the surrounding tissue, such algorithms are not feasible for brain extraction in this population, and fail to accurately segment the brain from tissues such as the placenta and the mother’s organs. The lack of brain extraction tools creates a major roadblock for fetal fMRI analysis.

In the past, brain extraction of fetal fMRI data was performed by manually (Thomason et al., 2014, 2015) or semi-automatically (Thomason et al., 2013; van den Heuvel et al., 2018) segmenting the brain from the surrounding tissue. As fMRI data generally consists of many acquisitions of multi-slice, 3D volumes, manual and semi-automatic segmentation can take over 30 hours per scan, a time-cost that is incredibly prohibitive for fetal researchers. Recently, Rutherford et al. (Rutherford et al., 2021) demonstrated the feasibility of an automated approach, using manual whole-brain segmentations to train a U-net convolutional neural network (CNN). Although this approach provides significant time-cost savings, it uses 2D convolutions on 2D slices, potentially hindering performance and generalization to new datasets. For example, Dice similarity coefficients of the overlap between manual segmentations and automated brain masks correlated significantly with gestational age, performing better in older fetuses. Thus, there remains a need for robust masking tools that perform well on fMRI data obtained in a range of participants.

Here, we present *funcmasker-flex*, a new BIDS App for masking fetal fMRI that overcomes the complexities of fetal fMRI brain masking with a robust 3D CNN architecture and is implemented in an extensible and transparent Snakemake workflow. Using locally acquired fetal fMRI data while also leveraging the large open dataset provided by Rutherford and colleagues, we provide an open-source fetal brain segmentation tool that performs well on data from a range of gestational ages and acquisition sites.

## 2. Results

### 2.1 Model performance

Performance of the baseline method (2D U-net) (1) and the proposed method (3D U-net) are shown in Fig. 1, comparing the distribution of Dice metrics on the training and generalization datasets. funcmasker-flex segmentations were consistently robust (all Dice metrics ≥0.74), while the baseline method produced segmentations with substantial errors (Dice < 0.6) in 4% of the source dataset volumes, and in 11% (238 volumes) of the generalization dataset volumes. In un-labelled data, visual quality control similarly demonstrated no observable errors in the funcmasker-flex outputs.

**Fig. 1.**
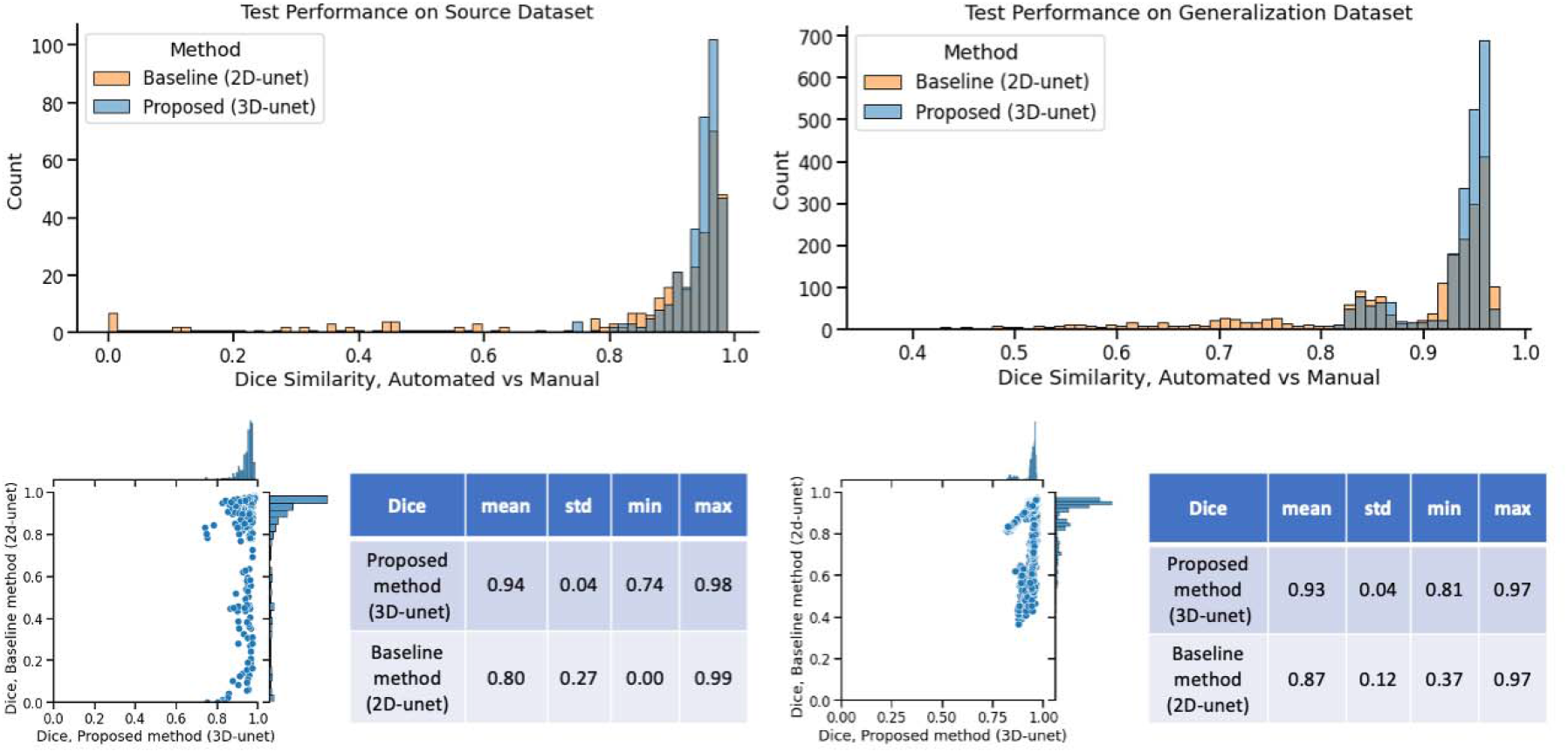
Comparison between the baseline 2D U-net model and the proposed 3D U-net model. Left, Dice similarity coefficients between manual masks and model performance when tested on the testing data of the source dataset, as well as distribution of Dice scores and descriptive statistics. Right, Dice similarity coefficients between manual masks and model performance when tested on the generalization dataset, as well as distribution of Dice scores and descriptive statistics.

### 2.2 Comparison to 2D CNN model

When comparing performance on the test dataset, *funcmasker-flex* performed significantly better than the baseline 2D U-net model, *V* = 28,882, *p* < .001. When comparing performance on the generalization dataset, *funcmasker-flex* again performed significantly better than the baseline 2D U-net model, *V* = 705,528, *p* < .001.

### 2.3 Performance based on scanner strength (1.5T versus 3T)

Dice similarity coefficients by scanner strength for each model are shown in Fig. 2. Linear mixed effects analysis showed a significant main effect of model (*F*(4,890.2) = 749.43, *p* <.001) and a significant model X scanner strength interaction (*F*(4,890.2) = 300.96, *p* < .001). Estimated marginal means revealed that at each scanner strength, the proposed 3D U-net model performed significantly better than the baseline model (all *ps* < .001), whereas there was no statistical difference between the 1.5T and 3T scanners within or between models (all *ps* > .05).

**Fig. 2.**
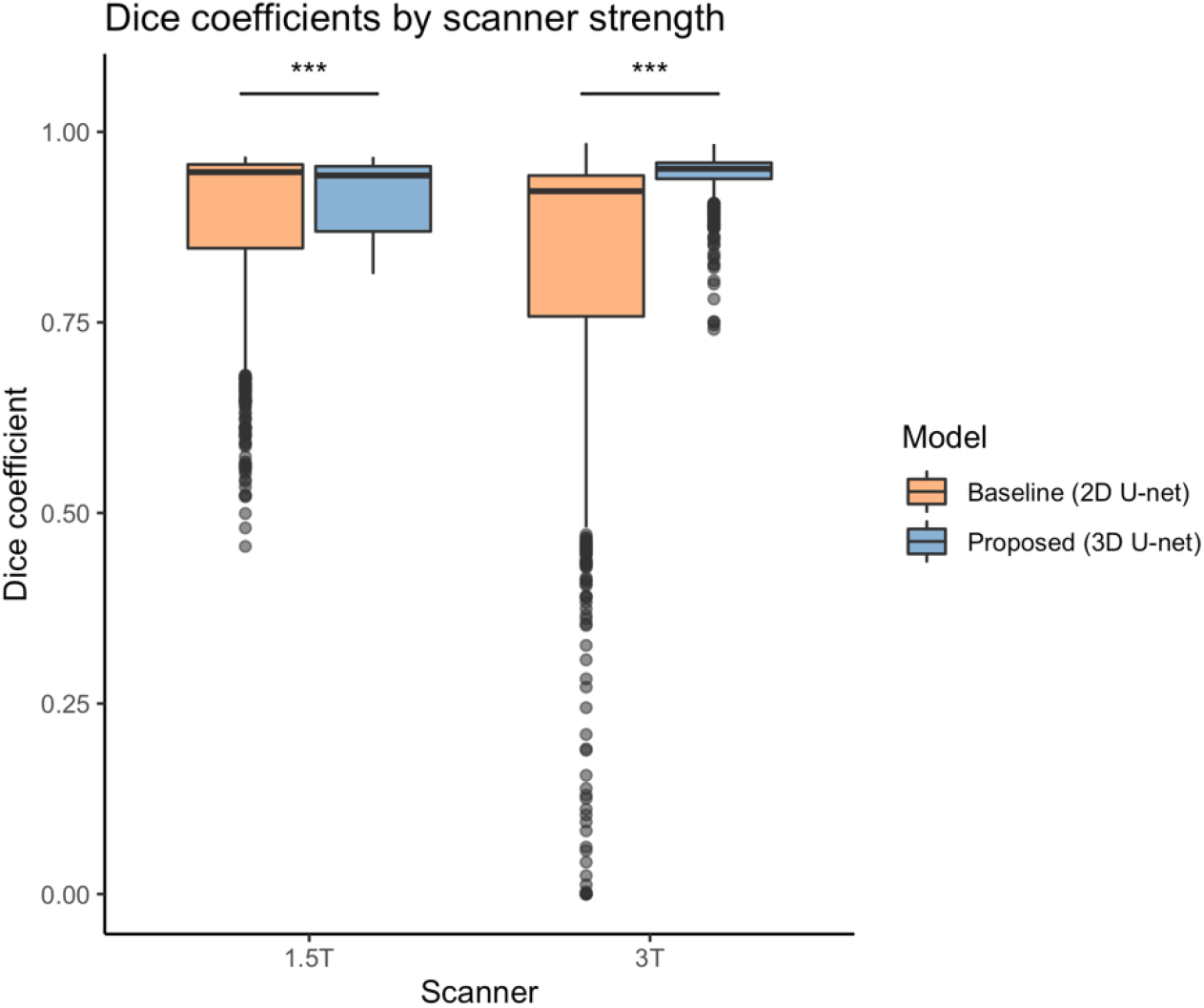
Dice coefficients by scanner strength for baseline and proposed models.

### 2.4 Correlation between Dice similarity coefficients and gestational age

Linear mixed effects analysis showed a significant main effect of model (*F*(4,890.1) = 57.84, *p* < .001) and a significant model X scanner strength interaction (*F*(4,890.1) = 33.31, *p* < .001). As shown in Fig. 3, there was no relationship between gestational age and dice coefficients in the proposed 3D U-net model, whereas the baseline 2D U-net model showed a positive relationship, with Dice similarity coefficients being higher at later gestational ages.

**Fig. 3.**
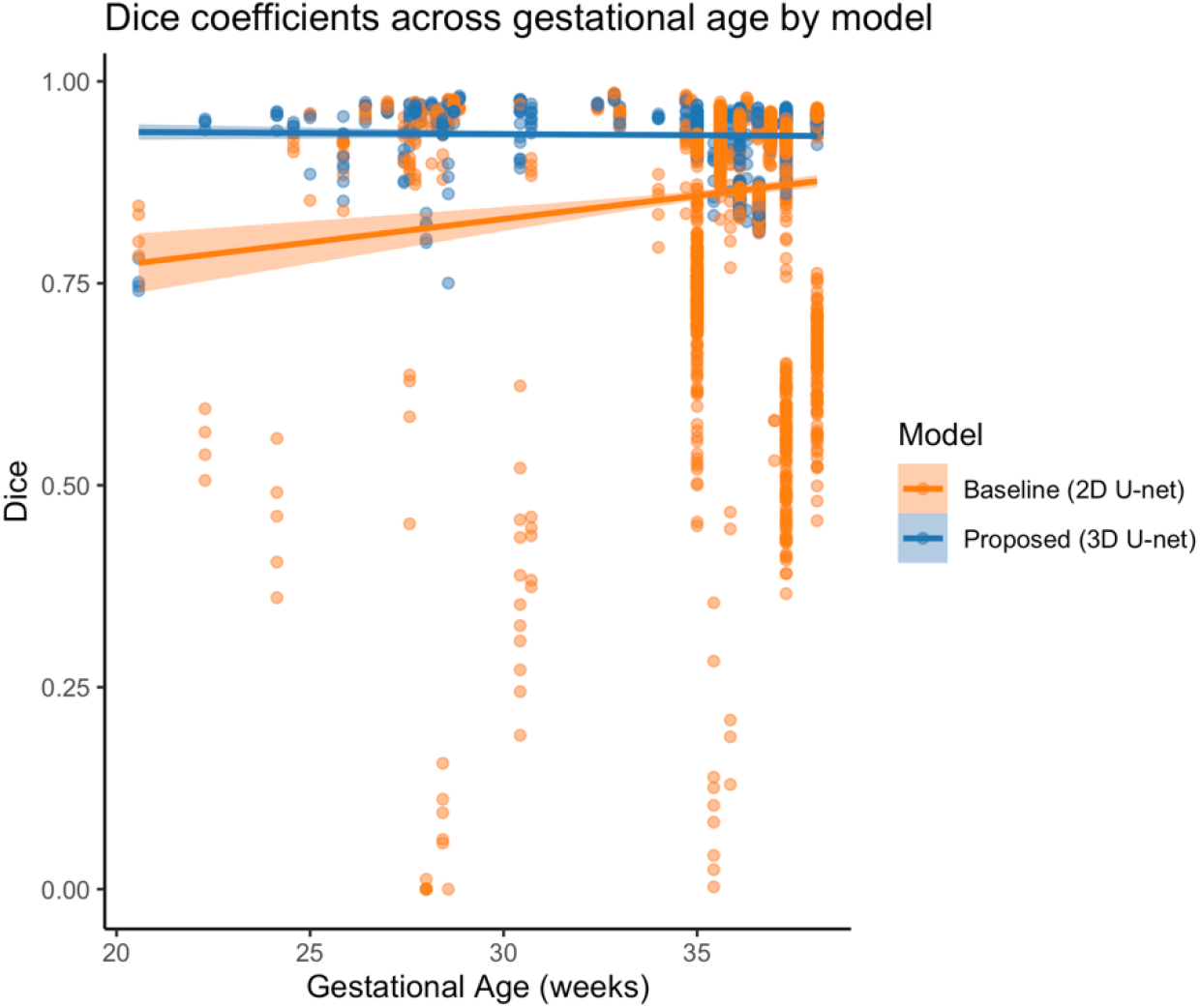
Dice coefficients by gestational age for baseline and proposed models.

## 3. Discussion

In this work, we present *funcmasker-flex*, an automated brain masking tool for fetal fMRI. Because standard brain extraction tools (e.g., BET by FSL) are not capable of delineating the fetal brain within the surrounding tissue, conducting analyses on fetal fMRI, in general, resulted in spending hundreds of hours manually segmenting the brain in each volume, a significant burden for researchers. To address this issue, we used a large set of manually-segmented fMRI volumes, we trained a 3D CNN to create a robust model to detect the brain within the surrounding tissue. Importantly, this model performed with high accuracy on an untrained dataset collected on a different scanner, demonstrating its generalizability.

When comparing *funcmasker-flex*’s performance on a new, untrained dataset to the gold standard manual tracing of the same data, we found high levels of similarity as measured by Dice similarity coefficients. That is, there was a high overlap in the spatial distribution of masks generated automatically by *funcmasker-flex* and the manual tracing. While manually segmenting the fetal brain in a single EPI volume took trained tracers approximately 17 minutes, automated segmentation took 1 minute and did not require a researcher to be present except to run the command, providing significant time-cost savings. A previous 2D CNN model (Rutherford et al., 2021) performed significantly less accurately on the untrained data, with Dice similarity coefficients ranging from .37 to .97. While this model still provides huge savings in terms of time spent manually segmenting the brain, it requires a large amount of manual correction to fill in missing segments of the mask.

We took several steps to make this tool easy to access, install, and use. First, the documentation and corresponding code is hosted on GitHub (https://github.com/khanlab/funcmasker-flex) and are open access. The pipeline was written in an extensible and transparent Snakemake workflow. It is also easy to install on any Linux machine with the command *“pip install funcmasker-flex”*, and will download any required dependencies, including containers (when the --use-singularity option is applied), when it is executed. It can be executed with a single command, and will work on any BIDS-formatted fetal fMRI dataset. If the user prefers not to download the entire package, it can also be run as a container, and example usage is provided in the documentation. The output also follows the BIDS naming convention, which means that it can easily be fed into other BIDS-dependent tools such as *fMRIPrep* (Esteban et al., 2019).

This study builds on previous work showing the feasibility of using CNNs in segmenting biomedical images (Isensee et al., 2021; Rutherford et al., 2021). Specifically, segmentation was performed using *nnU-Net*, a deep learning based method that automatically configures itself (Isensee et al., 2021). This framework showed a high degree of accuracy when segmenting several types of biomedical images, including the heart, liver, and kidneys. Building upon the *nnU-Net* framework allowed us to create a tool specific to the fetal brain, making use of open-source scientific tools.

We also build upon previous work showing the feasibility of CNNs in segmenting the fetal brain (Rutherford et al., 2021). This previous tool used a 2D CNN to mask 2D slices of fMRI data; while it performed well on some volumes in the untrained data, the overall Dice similarity coefficients of the overlap between manual and automated segmentations was low. Rutherford and colleagues showed a significant positive correlation with model performance and gestational age, suggesting that its generalizability may be limited to older fetuses. By using a 3D CNN, *funcmasker-flex* takes into account spatial boundaries between slices of a single EPI volume, improving performance as well as generalizability. Indeed, we did not see a significant correlation between model performance and age.

Although performance measures were generally high, there are several limitations to the tool that must be discussed. First, despite the robustness of *funcmasker-flex* in masking fetal brain volumes, visual inspection for quality control is still required. This is true however for all brain extraction methods; the output of tools such as FSL’s *BET* (Smith, 2002) and AFNI’s *3dSkullStrip* must be inspected for accuracy. It is also unclear how far the generalizability of *funcmasker-flex* can extend, for example to variations in strength of the scanner, although it performed equally well on data collected at 1.5T and 3T.

In summary, *funcmasker-flex* is a new tool that provides fast and robust brain masking of fetal fMRI data that requires no expertise, generalizes well, and will work on any fetal fMRI data in BIDS format. It is freely available and easy to use. It eliminates the burden of manual segmentation, and by reducing the time it takes to segment a fetal brain volume from roughly 37 hours to minutes, it removes a severe roadblock to performing fetal fMRI.

## 4. Materials and Methods

This research includes data from a cross-sectional study conducted at Western University as well as the openly-available dataset Rutherford et al. (Rutherford et al., 2021) (obtained from OpenNeuro, dataset identifier: ds003090). The Rutherford dataset (WS/YU) contains two cohorts, one from Wayne State University (WSU) and one from Yale University (YU). The study at Western University was approved by the Western University Research Ethics Board and all caregivers gave written informed consent. The WSU and YU studies were approved by the corresponding institutional ethics boards, and all caregivers gave written informed consent. An overview of the study cohorts is given in Table 1.

**Table 1.** Characteristics of each fetal cohort

| Cohort | Unique Individuals | Individuals with longitudinal data | Ages scanned (weeks gestational age) | Number of individual BOLD 4D scans | Number of 3D volumes per scan<br>$M \pm SD (min, max)$ | Analyses |
| --- | --- | --- | --- | --- | --- | --- |
| Western cohort | 8 | 0 | 33 – 38 | 19 | $110 \pm 0 (110, 110)$ | Primary |
| WS/YU cohort | 159 | 22 | 24 – 39 | 181 | $6.06 \pm 2.17 (2, 13)$ | Secondary |

### 4.1 Participants

#### 4.1.1 Western cohort

This cohort consisted of cross-sectional data from the third trimester. Between October 2018 and March 2020, 11 pregnant women were recruited for scanning at 33 – 38 weeks gestational age. Usable data were available for eight of the 11 participants. Inclusion criteria were singleton pregnancy and maternal age ≥18 years. Exclusion criteria were contraindication to safely undergoing non-contrast MRI, weight/body habitus that would prevent a successful MRI, suspected congenital anomalies, and concomitant substance use.

#### 4.1.2 WS/YU cohort

This cohort consisted of longitudinal data from the second and third trimester. Data from 159 fetuses were available, and 22 had second time points. Fetuses ranged in gestational age from 24 – 39 weeks. Inclusion criteria were singleton pregnancy, maternal age ≥18 years, no complications, and no contraindications for MRI.

### 4.2 Magnetic Resonance Imaging

#### 4.2.1 Western cohort

Fetal imaging data were acquired using either a 3T GE Discovery scanner (Milwaukee, Wisconsin, USA) and 32-channel torso coil at the Translational Imaging Research Facility (Robarts Research Institute, Western University, London, Canada; *n* = 5) or a 1.5T GE scanner with a GEM posterior and anterior array coil (London Health Sciences Center, London, Canada; *n* = 6). A minimum of one and a maximum of three blood oxygen level-dependent (BOLD) fMRI (TR: 2 s, TE: 45-60 ms [3T] / 60 ms [1.5T], flip angle 70°, voxel size 3.75×3.75×4 mm^3^, 22 slices, 110 volumes) were acquired in each fetus.

#### 4.2.2 WS/YU cohort

WSU fetal fMRI data were acquired on a 3T Siemens Verio scanner (Erlangen, Germany) using an abdominal 4-Channel Flex Coil. Functional images were acquired using an echo-planar sequence (TR = 2000 ms, TE = 30 ms, slice thickness = 4 mm, 360 volumes). Multi-echo resting-state sequences were also collected in a portion of these subjects (TR/ TEs: 2000/18,34,50). The Yale University cohort contains ten fetuses scanned twice longitudinally (gestational ages 30-36 weeks, M=32.7, SD=1.9). The YU data were acquired on a 3T Siemens Skyra scanner using a 32-channel abdominal coil (TR: 2 s, TE: 30 ms, slice thickness = 3 mm, 32 slices, 150 volumes). A subset of volumes from each acquisition were made available (Table 1).

### 4.3 Manual brain segmentation

For the Western cohort, manual brain segmentation of 2,090 volumes was performed on raw EPI scans using *FSLeyes* (McCarthy, 2021; Smith et al., 2004) by four tracers trained to identify the fetal brain, with each tracer segmenting an independent set of brains.

Segmentation took approximately 17 minutes per volume, and each scan had 110 volumes, leading to a total of 31.2 hours spent performing manual tracing per fetal fMRI scan. For the WS/YU cohort, manual brain segmentation was performed using BrainSuite software (Shattuck & Leahy, 2002).

### 4.4 U-Net architecture

Training and inference was carried out using nnU-net (Isensee et al., 2021), a framework that automatically configures a U-net network architecture for a new task based on basic dataset properties, and has been shown to outperform specialized pipelines in a range of segmentation tasks.

### 4.5 Model training

Workflows for training, testing, and evaluating fetal segmentation were built using Snakebids, a tool that allows Snakemake workflows to easily parse and create BIDS datasets and to function as BIDS Apps (Khan & Haast, 2021). Fetal fMRI data from the WS/YU cohort (1) was used for training and testing the nnU-net model, using the functional scans acquired using an echo-planar imaging (EPI) sequence from 112 fetuses for training, and 48 fetuses for testing. We used the same training and test splits as Rutherford et al, as these were made available in the code. The nnU-net 3D full-res model was trained using 5-fold cross-validation for 500 epochs using the automatically-configured parameters, and required approximately 24 hours for each fold when running on NVIDIA T4 GPUs. Inference does not require GPU-acceleration, and was implemented using models from all 5 folds, along with test-time augmentation, to provide a single prediction for each volume. Post-processing by nnU-net (including retention of the largest connected component) was disabled. The command-line interface, along with visualization of the participant-level (inference) workflow is shown in Fig. 4. The workflows can be applied to any BIDS dataset containing fetal EPI sequences and performs the appropriate resampling and padding for running inference, returning 4D binary masks for each EPI run in BIDS-Derivatives naming standards, and in the same space as the original bold datasets.

**Fig. 4.**
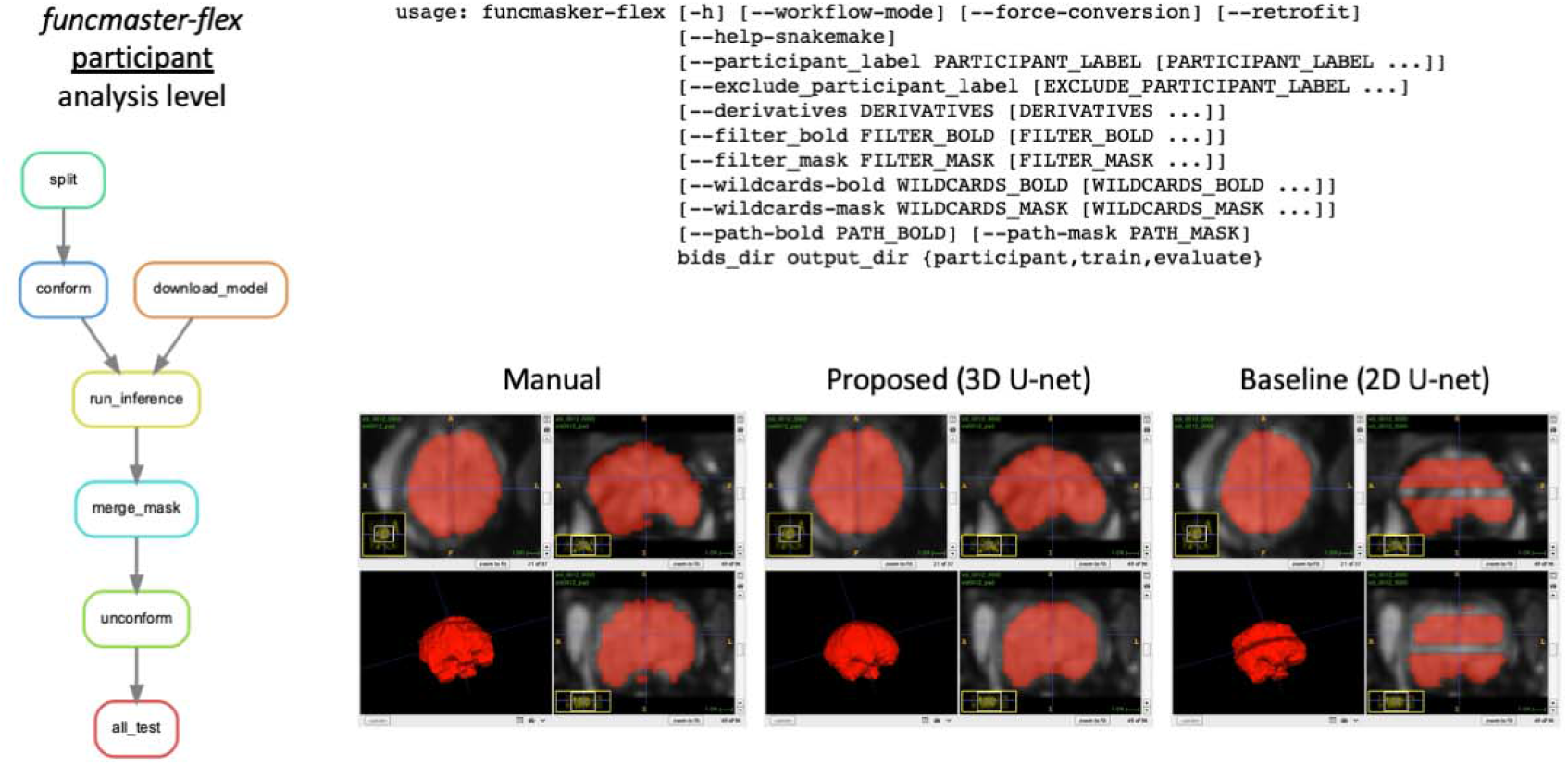
Left, visualization of the participant-level (inference) workflow. Top right, the command-line interface. Bottom right, example of a single volume mask created by manual tracing, the proposed 3D U-net model, and the baseline 2D U-net model.

### 4.6 Evaluation

We also tested generalizability of the model using 19 locally acquired functional scans from 8 fetuses (gestational age range=33-38 weeks, M=36.46, SD=0.98), which included 2,090 manually segmented 3D volume

### 4.7 Model comparison

Dice metrics were used to compare performance of the baseline 2D U-net approach (1) and the proposed 3D nnU-net approach as implemented in *funcmasker-flex* to the ground truth manually segmented volumes. A paired Wilcoxon signed-rank test was performed to determine whether the distributions of Dice similarity coefficients for each model statistically differed.

#### Performance based on scanner strength (1.5T versus 3T)

To determine whether scanner strength affected the accuracy of both the baseline 2D U-net and proposed 3D nnU-net approach, we examined Dice similarity coefficients across the two models. Linear mixed effects models were constructed with dice as the dependent variable, model (baseline/proposed) and scanner strength (1.5T/3T) as categorical variables and a model X scanner strength interaction. Participant was included as a random effect.

#### Correlation between Dice similarity coefficients and gestational age

To determine whether performance was affected by gestational age, we examined Dice similarity coefficients across ages in each model. Linear mixed effects models were constructed with dice as the dependent variable, model as a categorical factor, gestational age as a continuous factor, and a model X gestational age interaction. Participant was included as a random effect.

## Acknowledgements

The authors would like to thank the women who participated in these studies. We also thank the researchers at WS/YU for making their data and code available. We thank Megan Mueller, Sarah Abu Al-Saoud, Tajveer Ubhi, and Alissa Papadopolous for their assistance with manually tracing the fetal MRI data, and David Reese for his assistance in collecting the fetal MRI data.

## Funding

The funding for this research was provided by the Canadian Institutes of Health Research, the Molly Towell Perinatal Health Foundation and the Canada First Research Excellence Fund by BrainsCAN.

## Competing interests

The authors declare no competing interests.

## Notes

https://github.com/khanlab/funcmasker-flex

### Competing Interest Statement

The authors have declared no competing interest.

